# Adenosine triphosphate Binding Cassette subfamily C member 1 (ABCC1) overexpression reduces APP processing and increases alpha- versus beta-secretase activity, *in vitro*

**DOI:** 10.1101/2020.05.20.105064

**Authors:** Wayne M. Jepsen, Matthew De Both, Ashley L. Siniard, Keri Ramsey, Ignazio S. Piras, Marcus Naymik, Adrienne Henderson, Matthew J. Huentelman

**Author notes:** **Corresponding Author:** Matthew Huentelman, Ph.D., Professor, Neurogenomics Division, Translational Genomics Research Institute, 445 N Fifth Street, Phoenix, AZ 85004. **Contributions:** Wayne M. Jepsen designed, carried-out, and analyzed the experiments, as well as wrote the manuscript. Matthew De Both assisted with the analyses. Ashley L. Siniard contributed to the acquisition of data. Keri Ramsey contributed to the acquisition of data. Ignazio S. Piras assisted with the analyses. Marcus Naymik assisted with the analyses. Adrienne Henderson contributed to the acquisition of data. Matthew J. Huentelman is the principal investigator and guarantor of the work.

## Abstract

The organic anion transporter Adenosine triphosphate Binding Cassette subfamily C member 1 (*ABCC1*), also known as *MRP1*, has been demonstrated in murine models of Alzheimer’s disease (AD) to export amyloid beta (Abeta) from the endothelial cells of the blood-brain barrier to the periphery, and that pharmaceutical activation of ABCC1 can reduce amyloid plaque deposition in the brain. Here, we show that ABCC1 is not only capable of exporting Abeta from the cytoplasm of human cells, but also that it’s overexpression significantly reduces Abeta production and increases the ratio of alpha- versus beta-secretase mediated cleavage of the Amyloid Precursor Protein (APP), likely via indirect modulation of alpha-, beta-, and gamma-secretase activity.

## Background

Alzheimer’s disease (AD) is the sixth leading cause of death in the United States, and no current treatment exists that can effectively prevent or slow progression of the disease. For this reason, it is imperative to identify novel drug targets that can dramatically alter the physiological cascades that lead to neuronal cell death resulting in dementia and ultimately loss of life.

The deposition of aggregated amyloid beta (Abeta) in the brain is one of the major pathological hallmarks of AD, and Abeta species result from the differential cleavage of the Amyloid Precursor Protein (APP) (Selkoe and Hardy, 2016). APP is a single-pass transmembrane protein that is highly expressed in the brain and can be cleaved by a variety of secretases to produce unique peptide fragments, the two major pathways of which are known as the alpha- and beta-secretase pathways (Selkoe and Hardy, 2016). Cleavage by an alpha-secretase releases the soluble APP alpha (sAPPalpha) fragment from the membrane into the extracellular space, which has been shown to be neuroprotective and increase neurogenesis, in vitro (Ohsawa *et al.*, 1999), as well as play a positive role in synaptic plasticity (Ring *et al.*, 2007; Hick *et al.*, 2015) and memory formation (Bour *et al.*, 2004). Alpha-secretase cleavage of APP is the by far the most common cleavage of APP in the brain (Haass and Selkoe, 1993). If, instead, the APP molecule is cleaved by a beta-secretase, soluble APP beta (sAPPbeta) is released into the extracellular space, and subsequent cleavage of the remaining membrane-bound fragment by the gamma-secretase complex results in the production of Abeta, the peptide that aggregates to form amyloid plaques (Baranello *et al.*, 2015). Because alpha-secretases cleave APP within the Abeta domain, and beta-secretases cleave within the sAPPalpha domain, APP cleaved by an alpha-secretase cannot be cleaved by a beta-secretase, and vice versa (Haass and Selkoe, 1993), the two pathways are mutually exclusive and stoichiometrically related. It has been hypothesized that decreasing Abeta production could slow progression of AD, and although direct beta-secretase inhibition has failed in clinical trials (Das and Yan, 2019), control of these pathways via pharmaceutical intervention may still prove to be a viable AD treatment.

Our lab investigated the role of *ABCC1* in AD because we identified rare *ABCC1* single nucleotide polymorphisms (SNPs) in a familial case of late-onset AD, and in an early-onset AD patient with no family history of AD. ABCC1 has been previously shown to export Abeta from the cerebral spinal fluid to the peripheral blood (Krohn *et al.*, 2011), and *Abcc1* knockout mouse models have increased cerebral amyloid plaque deposition and soluble Abeta (Krohn *et al.*, 2011, 2015). Though our final experiments demonstrated no significant difference between the human reference *ABCC1* and either of the mutant alleles, our study revealed that *ABCC1* overexpression results in a significant reduction in extracellular Abeta1-40, Abeta1-42, and sAPPbeta species, while increasing the ratio of alpha-to beta-secretase mediated cleavage of APP, likely via indirect transcriptional modulation of proteins involved in APP metabolism. Our results indicate that ABCC1 is a valid drug target for the treatment of AD because of its multimodal influence on Abeta deposition: via exportation of Abeta species, as well as modulation of APP processing away from the amyloidogenic pathway.

## Results and Discussion

Briefly, for the APP metabolite experiments, ABCC1-overexpressing cells, or empty vector control cells, were plated at 1.4e7 cells per well of a 6-well plate weekly, with daily media changes. On the fourth day, supernatant was harvested and clarified, and cells were lysed for RNA or protein extraction. All samples were stored at −80 °C until 3 weeks of experiments were assayed together. A more complete description of the experimental approach can be found in the Materials and Methods section.

In the first experiment, ABCC1-overexpressing cells had a 34.02% decrease in extracellular Abeta1-40 (t=11.184, df=4, p=3.64e-04) and a 32.85% decrease in extracellular Abeta1-42 (t=4.26, df=4, p=0.013). The second experiment saw a 33.08% decrease in Abeta1-40 (t=5.26, df=4, p=6.26e-03) and a 43.90% decrease in Abeta1-42 (t=3.37, df=4, p=0.028) (see **Figure 1**). These results were surprising because, as previously stated, ABCC1 has been shown to export Abeta from the cytoplasm to the extracellular space, and if Abeta is, in fact, a substrate for ABCC1, we would expect to see higher extracellular concentrations of Abeta species.

**Figure 1:**
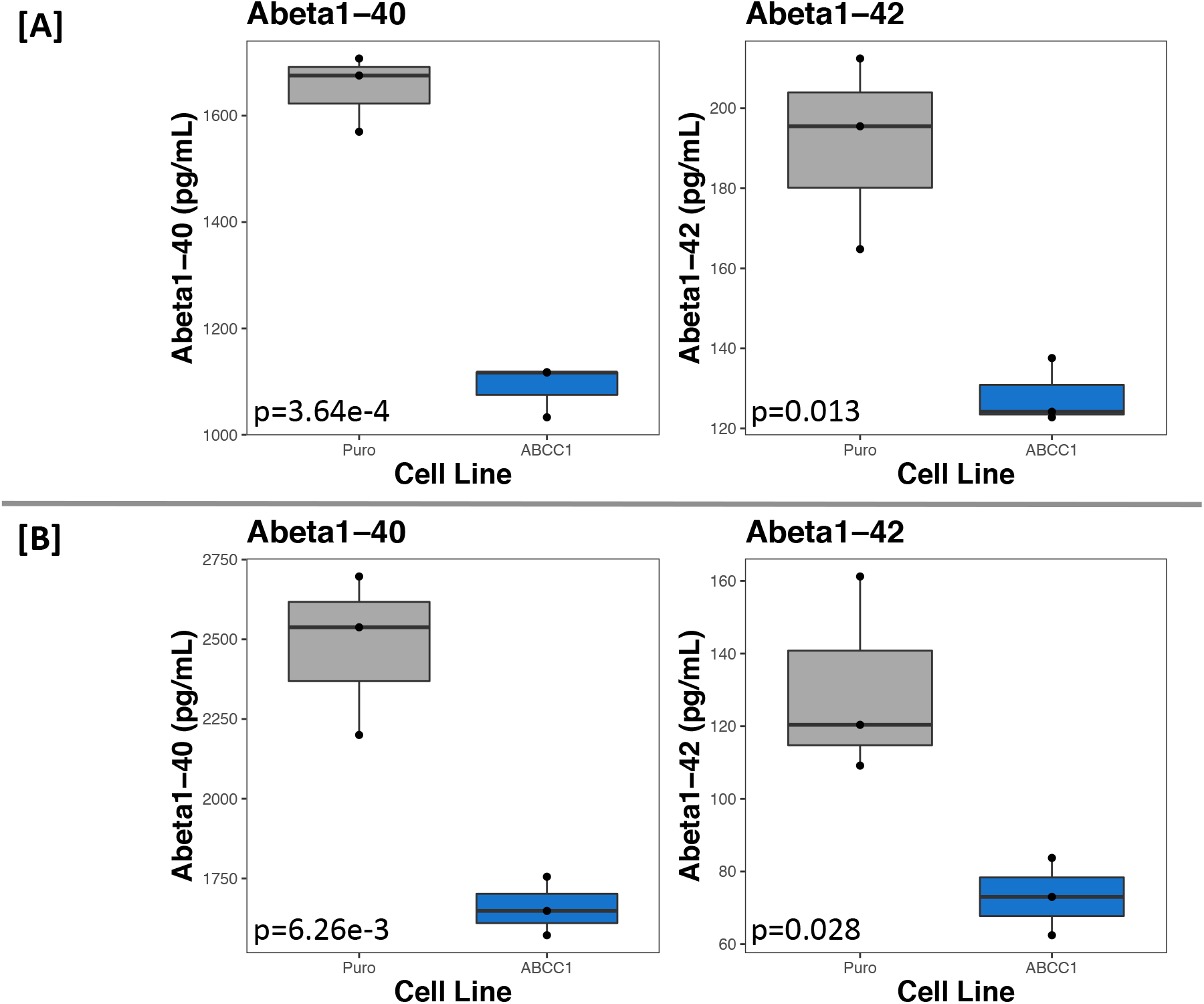
ABCC1 overexpression in BE(2)-m17 cells significantly decreases extracellular Abeta1-40 and 1-42 levels. [A] and [B] are experiment 1 and 2, respectively. The empty vector cell line is labeled “Puro” (grey boxes), and ABCC1-overexpressing cells are labeled “ABCC1” (blue boxes). Each point on the plots is the mean of technical quadruplicates, as measured by ELISA. P-values reported on each plot are calculated from Student’s two-sample t-test by comparing the two groups in that plot (N=6, n=3 for each plot).

To test whether ABCC1 exports Abeta, both cell lines were incubated with 200nM fluorescent Abeta1-42 (Beta-Amyloid (1-42), HiLyte Fluor 555-labeled, Human, AnaSpec, Fremont, CA, USA) for 18 hours, and then cells were subject to flow cytometry (FACSCanto II, BD Biosciences, Franklin Lakes, NJ) to quantify the percentage of fluorescent cells. 79.7% of the empty vector control cells were fluorescent, while only 68.4% of ABCC1-overexpressing cells displayed intracellular fluorescence. Furthermore, when incubated with fluorescent Abeta1-42 and 25uM thiethylperazine (MilliporeSigma, Burlington, MA, USA), a small molecule previously shown to increase ABCC1-mediated transport of Abeta (Krohn *et al.*, 2011), we observed a 23.6% decrease in population fluorescence in the empty vector control cells, and an even greater 38.0% decrease in the ABCC1-overexpressing cells (see **Figure 2**). This confirms that our model is working as expected because it agrees with previous reports: that ABCC1 does export Abeta, and that thiethylperazine increases ABCC1 transport activity.

**Figure 2:**
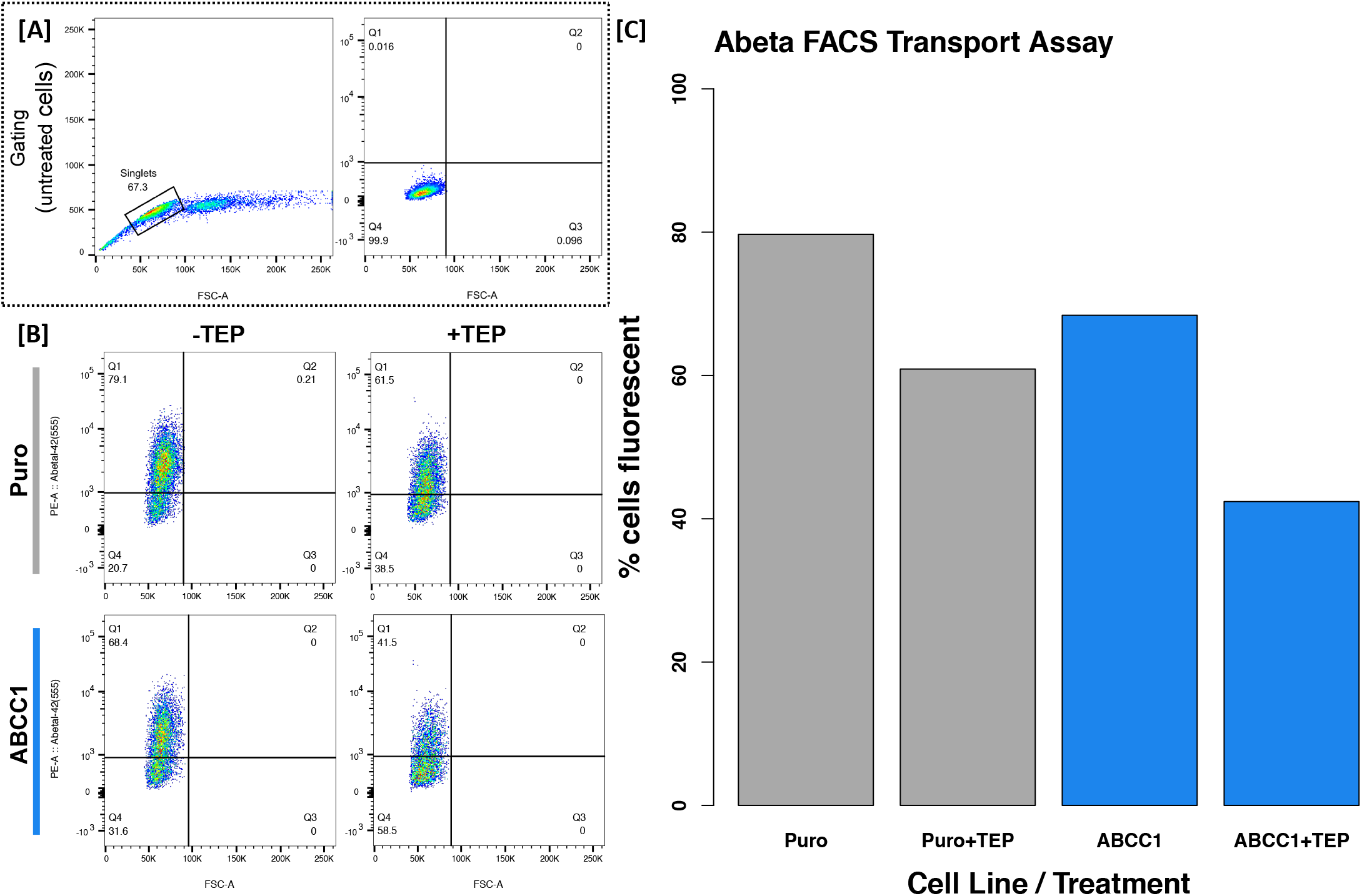
ABCC1 exports Abeta, and that activity is increased by thiethylperazine. [A] shows the original gating using untreated (unstained) cells to identify singlets and set a threshold for fluorescence. [B] shows the results of the cytometry experiment when empty vector cells (Puro, grey bars) or ABCC1 over-expressing cells (ABCC1, blue bars) are treated with fluorescent Abeta1-42 with or without thiethylperazine (TEP). Cells in quadrant 1 (Q1) are considered fluorescent, while those in Q4 are not. Percentage of fluorescent cells is plotted as a bar graph in [C].

Because we demonstrated that ABCC1 does export Abeta from the cytoplasm to the extracellular space, we hypothesized that ABCC1 may alter transcript levels of proteins capable of altering APP metabolism. To this end, we conducted RNA-sequencing of the cell lines. Analysis revealed 2470 differentially expressed genes (DEGs) with adjusted p-values less than or equal to 0.001, of which 2192 were protein coding. We hypothesized that because of the drastic reduction in extracellular Abeta, if a single gene were responsible for the altered APP processing, that it would have a log base two fold change (log2FC) with an absolute value greater than or equal to 1.5, which left 268 genes of interest (GOIs). Each gene was manually researched for their association to Alzheimer’s disease and amyloid pathology. This left 55 GOIs, 10 of which have known roles in APP/Abeta metabolism or transport – but whose expression levels are altered in the opposite direction one would expect for the observed ELISA results – and two with expression levels that may account for the lower levels of extracellular Abeta. All GOIs are discussed in **Supplementary Table 1** with a focus on this experiment, and in the context of the proceeding two RNA-seq experiments discussed later.

The genes whose expression levels may account for the reduced extracellular Abeta levels are *CD38* and *TIMP3*. *CD38* encodes the Cluster of Differentiation 38, an enzyme that synthesizes and hydrolyzes cyclic adenosine 5’-diphosphate-ribose, a molecule that regulates intracellular calcium signaling (Chini *et al.*, 2002). It has been shown that *Cd38* knockout AD mouse models have improved cognitive deficits, decreased cerebral amyloid burden, and that primary neurons cultured from those mice secrete significantly less Abeta species (Blacher *et al.*, 2015). The authors found that knockout of Cd38 alters beta- and gamma-secretase activity, effectively reducing both (Blacher *et al.*, 2015). This aligns with the observations made in our experiment, that when *CD38* expression is reduced (log2FC= −2.98, N=6, n=3, p=7.21e-09, padj=1.78e-07), extracellular Abeta levels are also reduced. Therefore, the reduction of *CD38* expression may contribute to the altered APP processing, though the mechanism by which ABCC1 alters *CD38* expression is not known.

*TIMP3*, our second candidate gene, encodes the Tissue Inhibitor of Metalloproteinases 3, a protein that can irreversibly inhibit APP-cleaving alpha-secretases like ADAM10 and ADAM17 (Hoe *et al.*, 2007). *TIMP3* expression is also reduced in AD brain tissue (Dunckley *et al.*, 2006), which may play a role in increased Abeta production. In our experiment, we saw TIMP3 expression reduced with a log2FC of −1.95 in the ABCC1-overexpressing cell line compared to the empty vector control (N=6, n=3, p=2.54e-110, padj=7.56e-107). Logically, if an alpha-secretase inhibitor is significantly decrease in expression, alpha-secretase activity would be increased, which would result in the reduction of secreted Abeta species because of the mutual exclusivity of the alpha-versus beta-secretase cleavage of APP previously discussed. It is also possible that the reduction of *CD38* and *TIMP3* works synergistically to reduce extracellular Abeta by decreasing beta- and gamma-, and increasing alpha-secretase activity.

To confirm our results, experiments were repeated with cryogenically preserved cells, as well as freshly transfected cells (to ensure that the transcriptional changes observed are not due to locus-specific integration of the transposable vectors), and APP metabolites were measured using the Meso Scale Discovery (MSD) platform (Meso Scale Diagnostics LLC, Rockville, MD, USA) which allows for the simultaneous, single-well measurement of Abeta1-40 and Abeta1-42, or sAPPalpha and sAPPbeta. Again, ABCC1-overexpressing cells had a 36.96% (t=10.97, df=10, p=6.74e-07) and a 35.21% (t=7.84, df=10, p=1.40e-05) reduction in extracellular Abeta1-40, as well as a 39.66% (t=11.42, df=10, p=4.66e-07) and 35.75% (t=9.73, df=10, p=9.03e-03) reduction in extracellular Abeta1-42. Furthermore, the two experiments saw a 29.45% (t=6.64, df=10, p=5.81e-05) and a 23.55% reduction (t=3.64, df=10, p=4.56e-03) in extracellular sAPPbeta, with no significant effect on sAPPalpha levels in the first experiment, but with a 16.27% reduction (t=3.21, df=10, p=9.30e-03) in the second experiment. Because the MSD platform allows for the simultaneous measurement of sAPPalpha and sAPPbeta in a single well, we used the ratio of sAPPalpha over sAPPbeta (sAPPalpha/sAPPbeta) to monitor alpha-versus beta-secretase cleavage of APP molecules because it controls for many of the confounding factors that could influence our measurements, and instead offers a mole-to-mole comparison. Indeed, in both experiments, ABCC1-overexpressing cells had a 35.20% (t=−10.89, df=10, p=7.24e-07) and 9.41% increase (t=−2.71, df=10, p=0.022) in sAPPalpha/sAPPbeta, implying a significant increase or reduction of alpha- or beta-secretase activity, respectively. Results are summarized in **Figure 3**.

**Figure 3:**
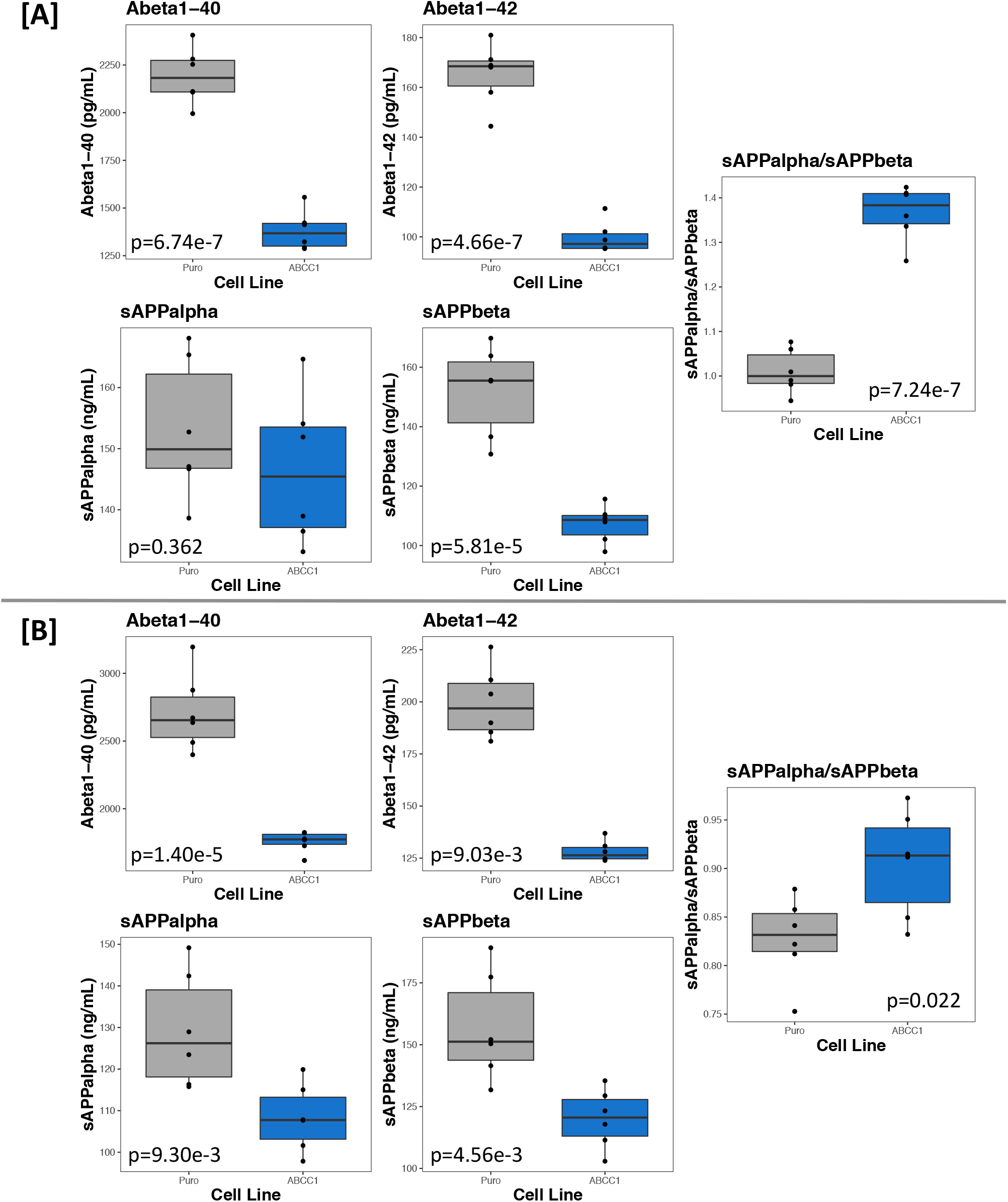
ABCC1 overexpression in BE(2)-m17 cells significantly decreases extracellular Abeta1-40, Abeta1-42, and sAPPbeta levels, and increases the ratio of alpha-to beta-secretase cleaved APP molecules. [A] and [B] show the third (cryopreserved cells) and fourth (newly transfected cells) APP metabolite experiments, respectively, measured using the MSD platform. The empty vector cell line is labeled “Puro” (grey boxes), and ABCC1-overexpressing cells are labeled “ABCC1” (blue boxes). All points on the plot are means of technical quadruplicates. P-values reported on each plot are calculated from Student’s two-sample t-test by comparing the two groups in that plot (N=12, n=6 for each plot). The results in [A] demonstrate that the decrease in extracellular Abeta species is not temporal, and [B] demonstrates that the location of integration of the transposable vectors is not the reason for altered APP metabolism.

Cells were again subject to RNA-seq. In both experiments, TIMP3 was significantly downregulated, with a log2FC of −0.64 in cryopreserved cells (N=6, n=3, p=0.015) and −0.82 in newly transfected cells (N=6, n=3, p=5.7e-03). CD38 had an insignificant log2FC of −0.53 in cryopreserved cells (N=6, n=3, p=0.24) and −0.48 (N=6, n=3, p=0.069) in the newly generated cell line; however, we do not believe that this is necessarily a reason to completely disregard the involvement of CD38 in the altered APP metabolism observed, as it is trending towards significance in the newly generated cell line. Furthermore, this confirms that the reduction in extracellular Abeta species is likely not due to integration of the transposable vectors within genes that alter APP processing, but rather that the increase in ABCC1 protein expression is likely altering transcription of genes whose products are capable of altering APP metabolism.

To determine if the transcriptional effects were cell line specific, we co-transfected the vectors (with SB100X) into ReNcell VM cells (MilliporeSigma), a human neural progenitor line, and extracted RNA from differentiated cells (14 days without growth factors). Transcripts were quantified using TaqMan (Applied Biosystems, Foster City, CA, USA) quantitative reverse transcriptase PCR (qRT-PCR), with targeted transcripts normalized *ACTB* expression, using the relative quantification (RQ) method (Livak and Schmittgen, 2001). *TIMP3* and *CD38* mean RQs were 11.10% lower (t=3.236, df=22, p=3.80e-03) and 76.0% lower (t= −12.76, df=22, p=1.21e-11), respectively, in the *ABCC1*-overexpressing cells versus the empty vector control (see **Figure 4**). These results agree with our previous results, that ABCC1 overexpression significantly alters the transcription levels of *TIMP3* and *CD38*, in a direction consistent with the reduced extracellular Abeta, and increased alpha-over beta-secretase cleaved APP molecules, and further demonstrates that altered transcriptional regulation of this gene is due to increased expression of ABCC1, rather than disruption of these genes due to transposable integration of the vectors.

**Figure 4:**
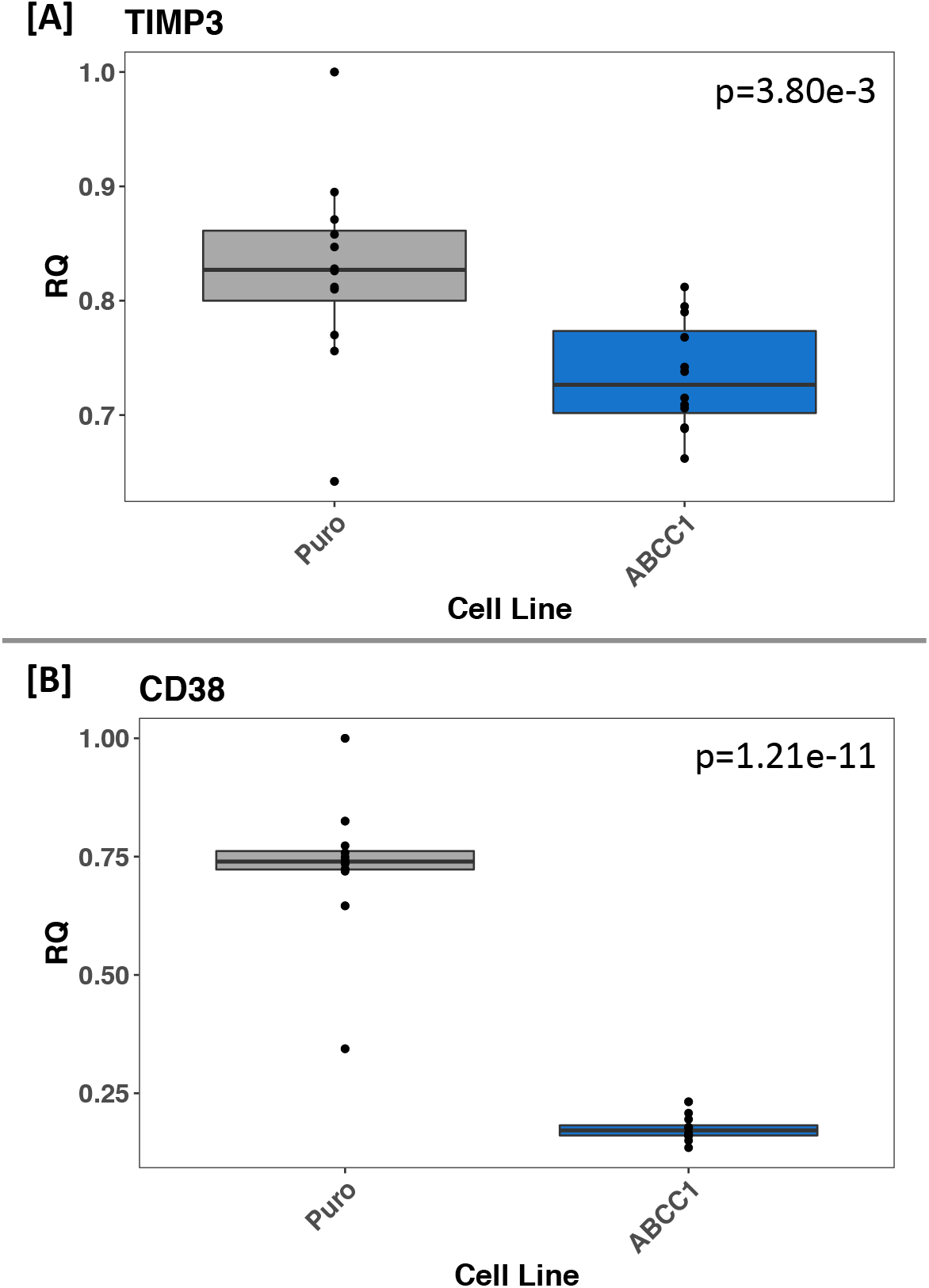
ABCC1 overexpression in ReNcell VM, a human neural progenitor cell line, significantly decreases mRNA levels of *TIMP3* and *CD38*. [A] and [B] show the relative quantification (RQ) of *TIMP3* and *CD38* mRNA in ReNcell VM. The empty vector cell line is labeled “Puro” (grey boxes), and ABCC1-overexpressing cells are labeled “ABCC1” (blue boxes). All points on the plots are means of technical quadruplicates. P-values reported on each plot are calculated from Student’s two-sample t-test by comparing the two groups in that plot (N=24, n=12 for each plot). The results confirm that altered expression of these genes is not due to location-specific genomic integration of the vectors, but rather because of increased ABCC1 expression. Furthermore, this demonstrates that decreased expression of these genes due to ABCC1 overexpression is not specific to the BE(2)-m17 human neuroblastoma cell line.

Taken together, our work confirms what previous labs have reported – that Abeta is a substrate for ABCC1-mediated export – but also provides novel insight: that increased ABCC1 expression reduces extracellular Abeta levels, likely via the alteration of alpha-, beta-, and gamma-secretase activity due to transcriptional modification of *TIMP3* and *CD38*. How ABCC1 alters these transcripts is unknown, but we hypothesize it is due to increased export of ABCC1’s canonical substrates, though further functional studies will be required to completely delineate the mechanism. Regardless, compounds that can dramatically increase ABCC1 transport activity or those that can increase ABCC1 expression, may prove to be viable drugs for the treatment or prevention of AD by not only increasing clearance of Abeta from the brain, but also by reducing the amount of Abeta that is produced. Many drugs have already been developed to block ABCC1 transport to prevent chemoresistance in cancer (Stefan and Wiese, 2019). Compounds identified in these drug development pipelines that have the opposite effect should be studied in the context of Alzheimer’s disease.

## Materials and Methods

### Cell line generation

#### BE(2)-m17

Human APP and ABCC1 codon-optimized cDNA was cloned into the Sleeping Beauty transposable vectors pSBbi-Hyg and pSBbi-Pur, respectively (gifts of Eric Kowarz, Addgene plasmids #60524 and #60523) by GenScript (Piscataway, NJ, USA). pSBbi-Hyg-APP was cotransfected with the transposase-encoding vector pCMV(CAT)T7-SB100 (a gift of Zsuzsanna Izsvak, Addgene plasmid #34879) into BE(2)-m17 human neuroblastoma cells (ATCC, Manassas, VA, USA), using the Cell Line Nucelofector Kit V and the Amaxa Nucleofector II Device (Lonza Group AG, Basel, CH). Stable cells were selected for with 1mg/mL hygromycin B (Invitrogen, Carlsbad, CA, USA). This APP-overexpressing cell line (now referred to as BE(2)-m17-APP) was then used to create the two experimental cell lines to ensure that APP expression is not variable due to transfection conditions. To this end, pSBbi-Pur-ABCC1 or empty vector was cotransfected with SB100X into BE(2)-m17-APP, and stable cells selected for with 10ug/mL puromycin (Gibco, Thermo Fisher Scientific, Waltham, MA), and maintained with 2ug/mL puromycin and 200ug/mL hygromycin B.

#### ReNcell VM

pSBbi-Pur-ABCC1 or empty vector were cotransfected with SB100X using the same Amaxa Nucleofector II and kit V (Lonza Group AG). Stably expressing cells were selected for using 10ug/mL puromycin (Gibco). When maintained with human epidermal growth factor (EGF) and fibroblast growth factor basic (bFGF) proteins (SigmaMillipore), ReNcell VM’s remain as human neuronal precursor cells. Upon removal of the growth factors, the cells will terminally differentiate and begin to mature into neurons and astrocytes.

### APP metabolite experiments and cellular extracts

Weekly, 1.4e7 cells per line were plated in a well of a 6-well plate without antibiotics (hygromycin and puromycin) and with daily media changes. On the fourth day, supernatant was harvested and supplemented to a final concentration of 1.0mM of an irreversible serine protease inhibitor, AEBSF (Thermo Fisher Scientific), then clarified at 10,000g for 10minutes at room temperature. Resulting supernatant was transferred to a new tube and stored at −80 °C until analysis. Cells in the plate were lysed for either protein extraction using RIPA buffer (Thermo Fisher Scientific) or RNA extraction using the Quick-RNA miniprep kit (Zymo Research, Irvine, CA, USA), either of which were stored at −80 °C until analysis.

For the first two sets of APP metabolite experiments, after 3 weeks of samples have been stored, supernatants were diluted 4-fold and assayed with the Amyloid beta 40 Human ELISA Kit and either the Amyloid beta 42 Human ELISA Kit or the Amyloid beta 42 Human ELISA Kit Ultrasensitive (Invitrogen), according to the manufacturer’s instructions. For the second two sets, supernatants were diluted 4-fold and assayed with the V-PLEX Plus Abeta Peptide Panel 1 (6E10) Kit and the sAPPalpha/sAPPbeta Kit (Meso Scale Discovery), according to the manufacturer’s protocol.

### Flow cytometry assay

Both cell lines were incubated with media supplemented with 200nM human Beta-Amyloid (1-42) HiLyte Fluor 555 (AnaSpec, Fremont, CA, USA), with or without thiethylperazine (MilliporeSigma) for approximately 18 hours. Cells were then washed twice with phosphate buffered saline (PBS), trypsinized, and spun-down. Pelleted cells were washed once with ice cold PBS, then resuspended in 1% FBS in ice cold PBS, and kept on ice until assayed. Sorting occurred on the FACSCanto II (BD Biosciences), and initially gated using untreated cells. Values are reported as the percentage of fluorescent cells.

### RNA sequencing

The first RNA-seq experiment utilized the TruSeq RNA Library Prep Kit v2 on the NextSeq500 (Illumina), and results mapped to 37,703 unique Ensembl IDs. The mean total reads per sample was 58.0±15.1 million. The next two RNA-seq experiments utilized the SMARTer Stranded Total RNA-Seq Kit v2 – Pico Input Mammalian (Takara Bio Inc., Kusatsu, Shiga, JP), and were sequenced on the NovaSeq 6000 (Illumina). Results mapped to 54,723 and 55,109 unique Ensembl IDs, respectively. The mean total reads per sample was 73.2±14.6 and 50.6±7.6 million reads, respectively. FASTQs were generated with bcl2fastq v2.18 (Illumina). Reads were aligned with STAR v2.7.3a (Dobin and Gingeras, 2016) to generate BAM files, and differential expression analysis was accomplished using featureCounts from Subread package v2.0.0 (Liao, Smyth and Shi, 2014) and DeSeq2 v1.26.0 (Love, Huber and Anders, 2014).

### qRT-PCR

Reverse transcription (RT) and no-RT reactions were achieved using SuperScript IV VILO Master Mix (Thermo Fisher Scientific), and qPCR was performed using TaqMan Fast Advanced Master Mix (Applied Biosystems) and multiplexed with primer/probe set for ACTB (Hs01060665_g1, VIC-MGB) and either TIMP3 (Hs00165949_m1, FAM-MGB) or CD38 (Hs00120071_m1, FAM-MGB). Reactions were run on the QuantStudio 6 Flex Real-Time PCR System (Applied Biosciences), according to the manufacturer’s protocol. Samples were measured in quadruplicate an quantified using the RQ = 2^(−(delta delta CT)) method (Livak and Schmittgen, 2001), and values reported as means of those technical replicates.

---

**Supplementary Table 1:**
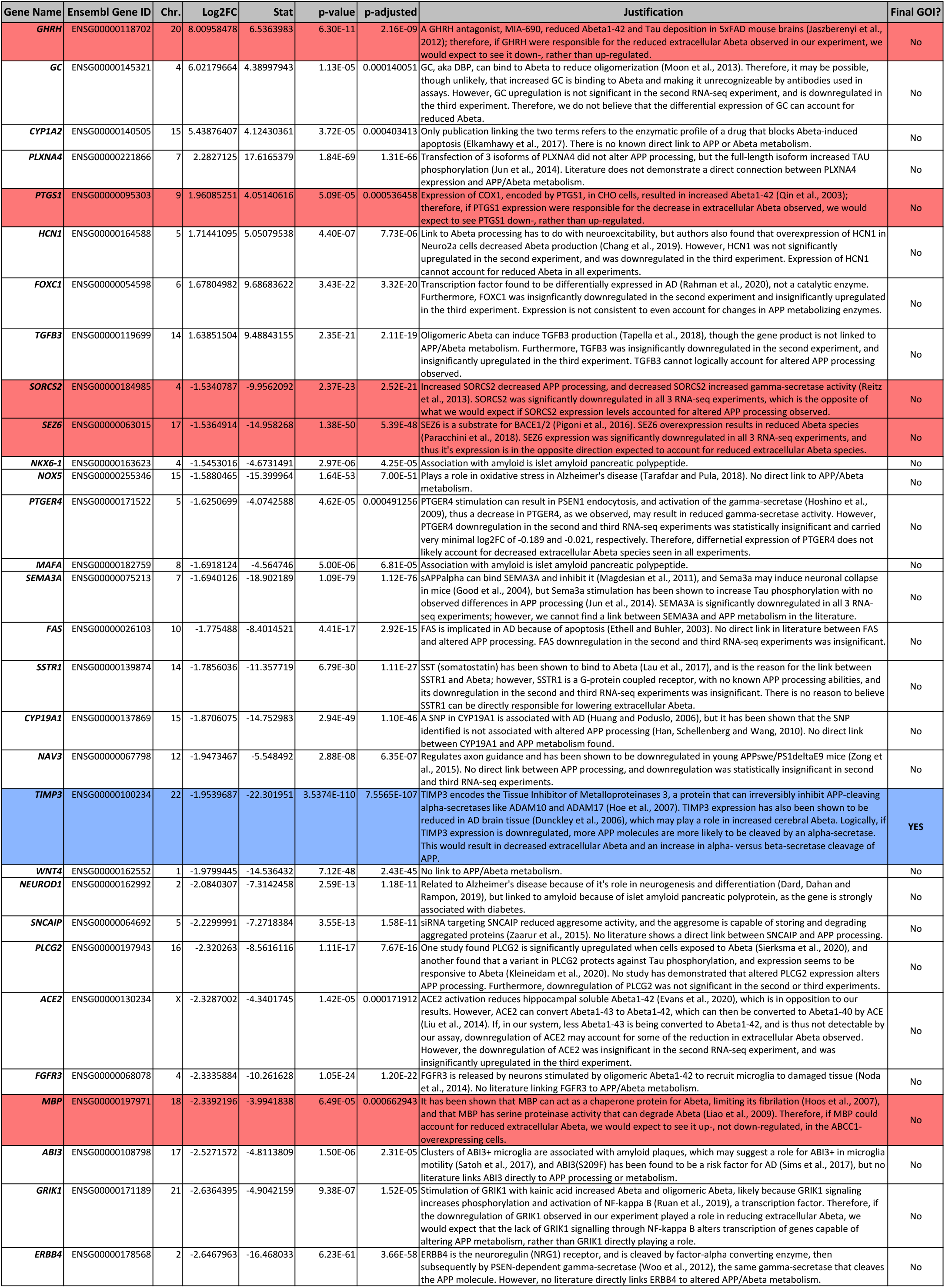
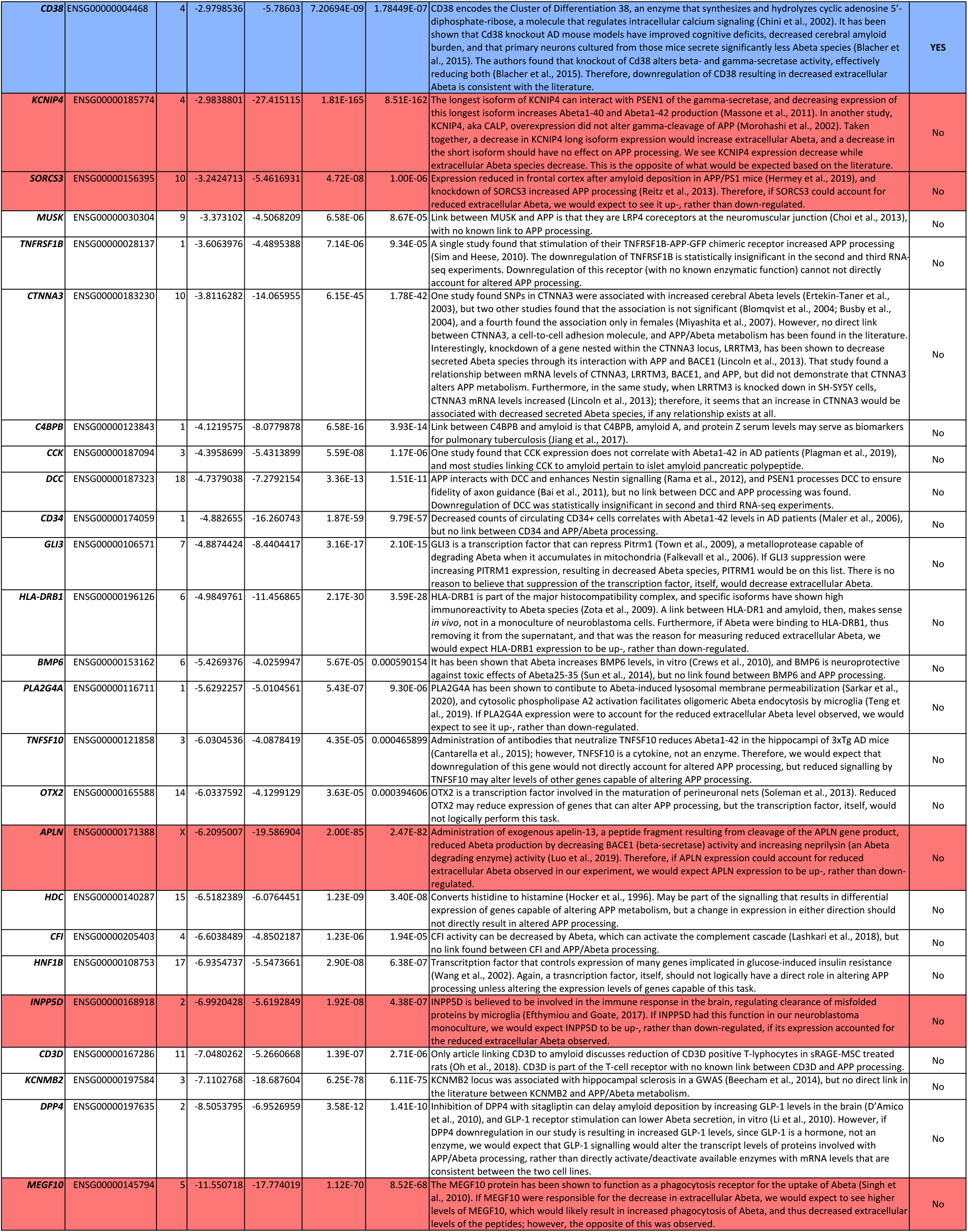
*TIMP3* and *CD38* are the only two differentially expressed genes with log2FCs that can account for the altered APP processing observed, according to current literature. Of the 55 GOIs identified, 12 are linked to altered APP processing or metabolism, but 10 of those genes have expression level changes in the opposite direction than are known in the literature to be able to account for the altered extracellular levels of APP-derived peptides observed. This may represent a lack of knowledge about these genes’ complete functions in APP metabolism, or it may indicate that their known roles have rather subtle effects. GOIs with expression levels consistent with our observations are highlighted in blue, while those that are directly inconsistent are highlighted in red.

